# Overground Gait Kinematics and Muscle Activation Patterns in the Yucatan Mini Pig

**DOI:** 10.1101/2021.10.19.465020

**Authors:** Soroush Mirkiani, David A. Roszko, Carly L. O’Sullivan, Pouria Faridi, David S. Hu, Daniel Fang, Dirk G. Everaert, Amirali Toossi, Kevin Robinson, Vivian K. Mushahwar

**Affiliations:** Neuroscience and Mental Health Institute, University of Alberta, Edmonton, AB, Canada; Division of Physical Medicine and Rehabilitation, Department of Medicine, University of Alberta, Edmonton, AB, Canada; School of Physical Therapy, Belmont University, Nashville, TN, United States of America; Sensory Motor Adaptive Rehabilitative Technology (SMART) Network, University of Alberta, Edmonton, AB, Canada

## Abstract

A growing number of spinal cord injury, neuromodulation, and cell therapy studies on porcine models, especially the Yucatan minipigs (YMPs), have been recently reported. This is due to the large similarities between human and porcine neuroanatomy and biomechanics. To assess treatment modalities and locomotor recovery in this model, there is an obvious need for detailed characterization of normative overground gait in neurologically intact YMPs. The objective of this study was to assess gait biomechanics and the effect of overground walking speed on gait parameters, kinematics, and electromyographic (EMG) activity in the hindlimb muscles of YMPs. Nine neurologically-intact adult YMPs were trained to walk overground in a straight line. Whole-body kinematics and EMG activity of hindlimb muscles were recorded and analyzed at 6 different speed ranges (0.4-0.59, 0.6-0.79, 0.8-0.99, 1.0-1.19, 1.2-1.39, and 1.4-1.6 m/s). A MATLAB program was developed to detect strides and gait events automatically from motion-captured data. Significant decreases in stride duration, stance and swing times and an increase in stride length were observed with increasing speed. A transition in gait pattern occurred at the 1.0m/s walking speed. Significant increases in the range of motion of the knee and ankle joints were observed at higher speeds. Also, the points of minimum and maximum knee and ankle joint angles occurred earlier in the gait cycle at higher speeds. The onset of EMG activity in the biceps femoris muscle occurred significantly earlier in the gait cycle with increasing speed. A comprehensive characterization of overground walking in neurologically-intact YMPs is provided. These normative measures set the basis against which the effects of future interventions on locomotor capacity in YMPs can be compared.

## Introduction

Neurological disorders including spinal cord injury (SCI) affect sensorimotor and autonomic function and impact the quality of life of millions around the world. In the United States, Hirtz et. al.[1] estimated that 4.5 in 100,000 people acquire a SCI annually. SCIs can cause alterations in locomotor, cardiovascular, respiratory, or urinary function and increase the risk for secondary complications such as pressure injuries[2]. To quantify the nuances of locomotor deficits and recovery, gait analysis has become a popular method for both humans and animals[3]. When paired with locomotor training after injury, gait analysis can be used to more precisely quantify the subtle, yet crucial, changes in gait parameters.

In developing treatments and devices for neurological disorders, pre-clinical studies in animal models are an important step to validate treatment efficacy and safety[4]. Rats and mice are common animal models in research; however, dimensional, physiological, pathological, and functional differences exist between rodents and humans[5], [6]. This underlies the critical need for large animal models in translational research[7]. Several studies have used feline[8], [9] canine[10], ovine[11], non-human primates (NHP)[12], and porcine models[6] in various experimental contexts. While pigs and cats are different than humans, the size of their spinal cords are more comparable to that of a human than rodents[6], [13]. NHPs, the most similar animals to humans genetically, can be the best model for behavioral or cognitive-based studies[14]. Also, the opposable thumbs in NHPs make them an ideal model for cervical SCI and finger movement studies. However, the access to primates and the ability to house them properly present ethical, technical and cost related problems to researchers, limiting the use of NHPs in translational experiments[15].

Several studies have used porcine models in SCI, traumatic brain injury and neuromodulation studies[6], [16], [17]. This is because the gross anatomy of the brain, vertebral column, spinal cord, and the ratio of cerebrospinal fluid volume to spinal cord volume in pigs have many similarities to that of humans[13]. Moreover, pigs are friendly, easier to handle than primates, and their size is well-suited for testing surgical setups and devices designed for humans. Therefore, they have been promising candidates for testing different neuromodulation strategies such as intraspinal microstimulation[7], [18], epidural stimulation,[19] and photo-biomodulation[20]. This is done using devices and surgical technologies identical to those performed intraoperatively, or intended to be performed, in humans[20], [21].

Domestic pigs gain weight non-linearly during their first two years of life and can reach more than 300 kg. On the other hand, miniature pig breeds with slower skeletal growth rates, such as: Yucatan, Panepinto, Hanfrod, Göttingen, and Vietnamese Pot-Bellied pigs, approach weights of 55-60 kg as adults, and have recently been widely used as animal models[6], [22], [23]. Although the porcine thoracic injury behavioral scale (PTIBS) and the Miami porcine walking scale (MPWS) have been widely used for gait analysis, they are based on visual observations and mainly qualitative assessments. Recently, Boakye et al.[22] characterized treadmill-based gait kinematics of Yucatan minipigs (YMPs). Several differences and similarities between treadmill-based and overground walking have been previously shown in terms of various spatiotemporal gait parameters, muscle activation, and joint kinematics[24]–[28]. Because the ultimate goal of treadmill-based training and neuromodulation paradigms is the translation to functional overground locomotion, there is a critical need for detailed characterization of overground walking parameters in pigs for studies focused on locomotor recovery. In this paper, we analyze overground gait kinematics of neurologically-intact adult YMPs using both motion capture and recorded EMG data. This information can be used as a standard guide to which future studies on injured minipigs undergoing various treatments can be compared.

## Methods

### Experimental Setup

All procedures were approved by the University of Alberta Animal Care and Use Committee. Nine adult female YMPs (weight=39.3±2.89 kg) were trained to traverse a straight 3.5 m walkway by following a human experimenter (Figure 1a, c). Twenty-seven reflective motion capture markers (Vicon Motion Systems Ltd, Oxford, UK) were mounted on the pigs to mark the locations of the left and right hindlimb hoofs, metatarsophalangeal (MTP) joints, ankles, knees, hips, iliac crests, forelimb hoofs, metacarpophalangeal (MCP) joints, wrists, elbows, shoulders, and scapulae, as well as the sacrum, mid-thoracic spine, and mid-cervical spine (Figure 1b). The positions of motion capture markers were recorded by eight cameras. Wireless surface EMG (sEMG) electrodes (Noraxon USA Inc., Scottsdale, AZ, USA) were placed on the left and right gluteus medius (GM) muscles, the biceps femoris (BF) muscles, and the vastus lateralis (VL) muscles (Figure 1b). For each pig, motion capture data (150 frames/sec) and EMG activity (3000 Hz) were recorded as the pigs traversed the walkway 20-120 times on a given testing day (average number of passes per day=65±28).

**Figure 1.**
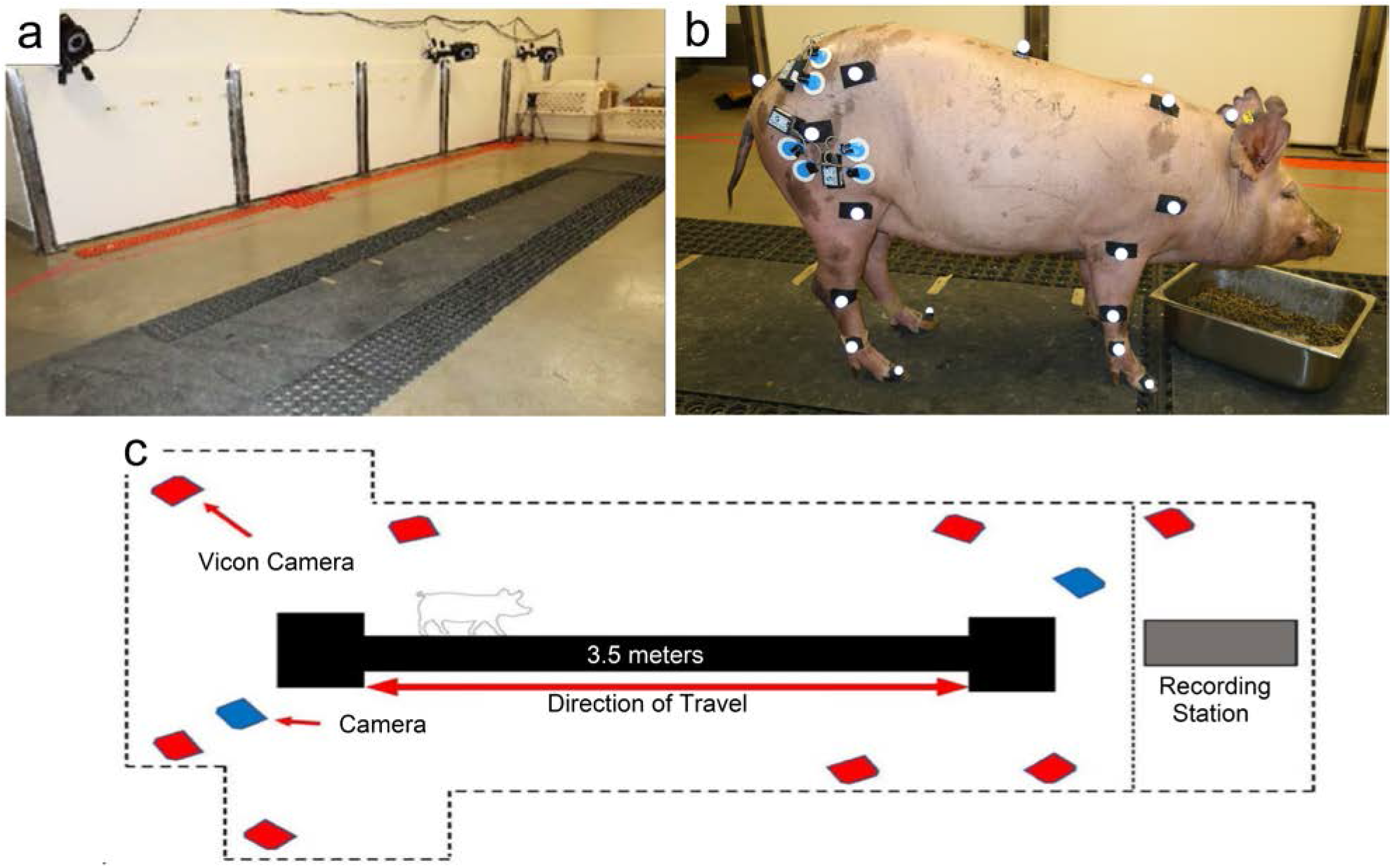
Experimental setup. An image **(a)** and schematic diagram **(c)** of the overground walkway used for characterizing YMP kinematics and EMG activity. **(b)** Position of motion capture markers and sEMG electrodes placed on a YMP.

### Automated Stride Detection

A MATLAB (R2020b, The MathWorks, Inc., Natick, MA, USA) program was developed to detect strides from raw motion capture data. Hoof contact (touch-down) and lift-off (toe-off) frames were detected by analyzing the kinematic data of the MTP markers for the hindlimbs or the MCP markers for the forelimbs. MTP/MCP markers were selected for detecting touch-down/toe-off events because the toe markers often became detached during the measurement sessions. During walking, when the limb is in contact with the ground, the y-values (along the horizontal direction of locomotion) remain almost unchanged until the limb is lifted and pushed forward again (Figure 2a). Therefore, by detecting the onset and endpoint of the duration when the y-values remain constant, the touch-down and toe-off frames can be detected. The y-coordinate was selected instead of the z-coordinate (vertical height of the marker) due to its uniform and repetitive trend, making it a reliable candidate for stride detection. Y-values of the MTP/MCP markers were extracted from the CSV files generated by the motion capture software and a zero-phase forward and reverse digital IIR filter (5 Hz lowpass) was applied to remove noise (Figure 2a). Because during each walkway pass y-values increase or decrease (during left-to-right or right-to-left walking, respectively), the y-value data corresponding to each pass were detrended (Figure 2b). A touch-down or toe-off event was then detected when the y-values of the MTP/MCP markers crossed constant, expert-defined thresholds. A sign function was used to determine when each touch-down or toe-off had occurred. For each motion captured frame, the difference between the magnitude of y-values of the MTP/MCP markers and one of the constant thresholds was fed into the sign function. The outputs of the sign function were compared with each other to detect appropriate touch-downs or toe-offs. The detected touch-down/toe-off frames were then classified to detect a stride. Each stride contained a touch-down, a toe-off and a next touch-down frame. Figure 2c shows that left-to-right and right-to-left runs on the walkway result in mirrored y-value trends. Because uniform minima were used, two different plots were needed: one for detecting touch-downs and one for detecting toe-offs. These two plots are inverse (negative) of each other.

**Figure 2.**
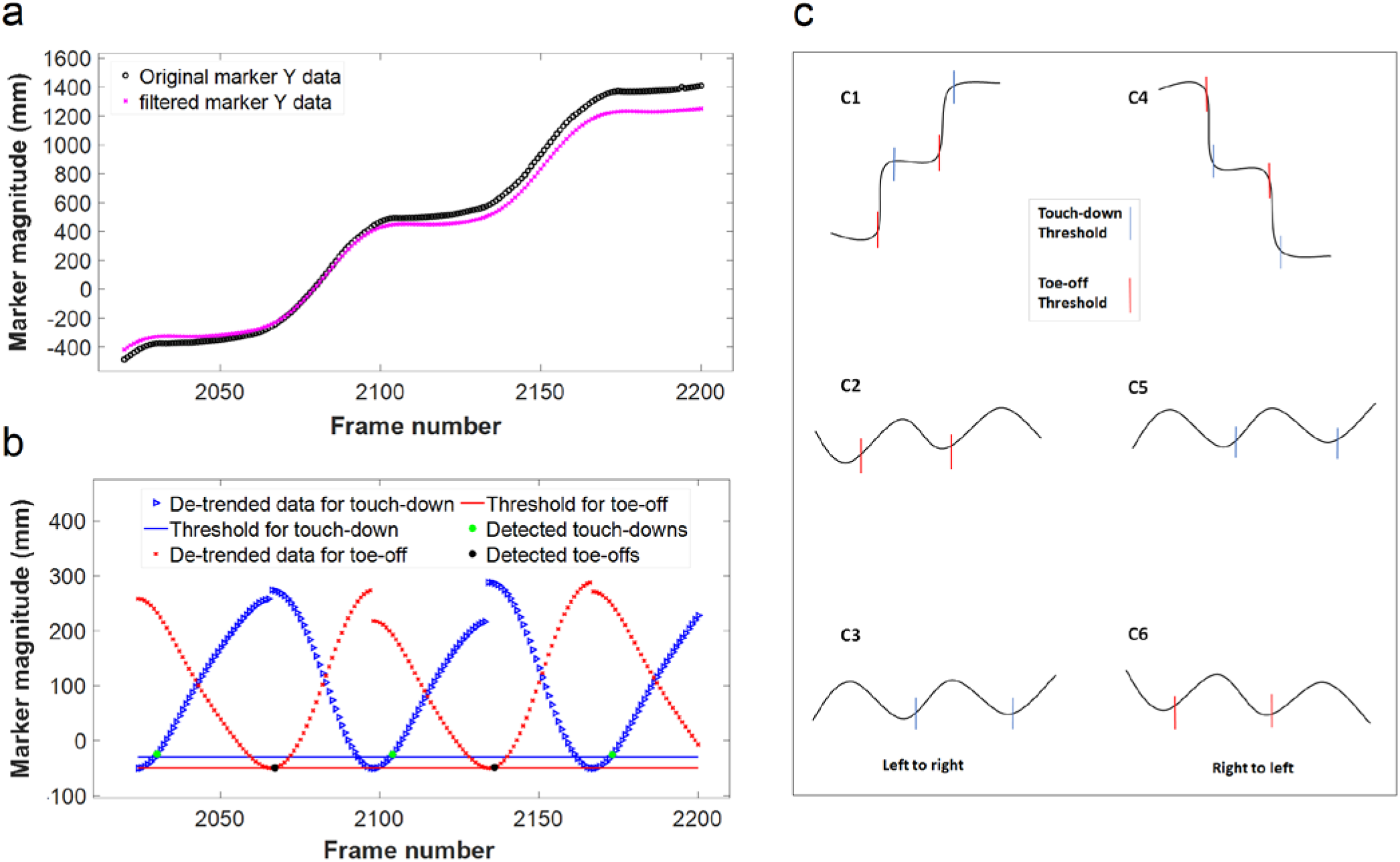
Y-values of a MCP joint reflective marker for walking from the left to the right end of the walkway. **(a)** Raw (○) and filtered (×) values. **(b)** Two detrended plots; one for detecting touch-down (▶) and one for detecting toe-off (×). Solid horizontal lines show the chosen thresholds for detection of the stride events. Once the marker y-value passed its preset threshold, its respective frame number was considered as a touch-down (green points) or a toe-off (black points). **(c)** An example of y-values in a back-and-forth movement across the walkway. In **(c1),** walking left to right, y-values had an incremental trend. After detrending, the plot was used for toe-off detection **(c2).** The plot was then flipped horizontally for touch-down detection **(c3).** The opposite picture can be seen on the for walking right to left **(c4)** where the y-values have a decremental trend. After detrending **(c5),** the plot was used for touch-down detection, and then horizontally flipped **(c6)** for toe-off detection. Note that 2 unique thresholds, one for touch-down and one for toe-off are used. The thresholds are depicted as horizontal lines in (b); threshold crossings are depicted by vertical lines (in c) for increased clarity.

As the pigs turned around at each end of the walkway highly variable data were produced. Therefore, the start and end frame of each section of straight walking was manually determined for the stride labelling program to work effectively. The program automatically detected the desired y-values of MTP/MCP markers for each walkway pass using two CSV files: one containing raw motion capture data and another containing the start and end frames of straight walking. The reconstructed strides were stored as an array in MATLAB and published as a CSV file for further analysis. Each file contained the detected strides of a specific limb for all walkway passes from one recording day. The accuracy of the program was validated by three experimenters who manually detected the events in motion capture videos. The program was deemed to be accurate if the automatically detected stride events were within 4 frames (26.6 ms) of the manually detected events. Across the three experimenters, the program was accurate 89.3±2.3 % (Mean±SD) of the time. The inaccurately detected events resulted in identified outliers in the kinematics data that were removed as explained in the next section.

### Analysis of Kinematic Data

Kinematic data were stored in CSV files and processed with custom-written MATLAB programs. The data were filtered with a zero-phase forward and reverse digital infinite impulse response (IIR) 10 Hz lowpass filter to remove vibration artifacts of the markers during walking. For each stride, stride speed, stride length, stance time, and swing time were calculated. Moreover, the height of the left and right iliac crests (pelvis), MTP height, hindlimb toe height, and ankle, knee and hip joint angles were calculated.

Because the hoof reflective markers were often lost during walking in five out of nine pigs, stride speed was calculated based on the left MTP marker. For each pig, strides were sorted into one of the 6 speed ranges, and within each speed range, the gait cycle was divided into 50 bins. The mean and standard deviation of the joint angles of the forelimbs and hindlimbs were then calculated for each bin. All data were plotted in reference to the left hindlimb, normalized to 100% of the gait cycle. Irregular outlier strides (caused by inaccurate event detection) for each pig were identified as strides that differed by greater than 3 standard deviations from the mean in the joint angle calculations and were removed. The mean joint angles across all 9 pigs were then averaged to provide an overall summary of the joint angles for YMPs during locomotion. If fewer than 6 strides were recorded for a given pig at a given speed, these data were excluded from the overall average calculation.

Interlimb coordination (coupling between limbs within a stride) was calculated as the fraction of overlap between steps of different limbs during the gait cycle[22]. Diagonal (LF-RH), homolateral (LF-LH), and homologous (LF-RF) coupling were calculated for strides within each of the 6 speed ranges. All coupling values were calculated with reference to the left forelimb and presented in polar plots to show interlimb phase relationships. Joint angle cyclograms were created for the left and right knee-ankle, hip-ankle, and hip-knee joints to investigate inter-joint coordination in the hindlimbs during overground walking. Cyclogram areas were calculated using the polyarea function in MATLAB. Regularity of cyclograms was calculated as a measure of repeatability of the steps using the approach described by Tepavac & Field-Fote[29]. Each regularity has a value between 0 and 1 that measures how repeatable a group of strides is. A score of 0 means that the cyclograms (strides) are nothing alike and a score of 1 means that the strides are identical[29].

### Analysis of EMG Activity

The EMG data were stored in CSV files and processed using a custom-written MATLAB program. The data were bandpass filtered (10 Hz - 450 Hz), full-wave rectified, and lowpass filtered at 10 Hz. All filtering was performed with zero-phase forward and reverse digital IIR filters to avoid phase distortion and delays. EMG activity was synchronized with the kinematic measurements using a common trigger in both the EMG and kinematic data. EMG data for the left and right biceps femoris (BF), vastus lateralis (VL) and gluteus medius (GM) muscles for each stride were sorted based on stride speed into one of the 6 defined speed ranges, and analyzed to determine the stride onset and offset points. EMG traces that were clipped (i.e., saturated the amplifier) along with traces with major artifacts, were manually removed. Outliers were removed using the same criteria applied to the kinematic data. If fewer than 6 strides were recorded for a given pig at a given speed, these data were excluded from further analysis.

A threshold was defined based on regions of consistently low muscle activity (mean + 1.5 SD) and EMG activity rising above this threshold was identified as an onset point while EMG activity dropping below this threshold was identified as an offset point [12]. Bursts that were separated by 20% or less of the gait cycle were combined and bursts with durations less than 10% of the gait cycle were then discarded. Finally, the onset and offset points of all strides exhibiting one burst of activity were averaged for each pig. These onset and offset values were then averaged across all pigs to determine the mean onset and offset points for all pigs for strides within each speed range.

### Statistical Analyses

Statistical analyses of joint angles, MTP and hind-hoof clearance above ground, the vertical left and right displacement of the pelvis and the difference between them (hip-hike), and EMG onset and offset points within the different speed ranges were performed using repeated measures analysis of variance (ANOVA; IBM SPSS, Build 1.0.0. 1447; IBM Corp., Armonk, N.Y., USA). Differences in stride lengths across the 6 speed ranges and between the four limbs within a speed range were analyzed using two-way repeated measures ANOVA. Interlimb and inter-joint coupling, swing/stance durations, and duty factor values (stance time divided by stride time) were compared across all speed ranges using one-way ANOVA. Quadratic, exponential, and power curve models were fitted to the swing time, stance time, and stride time distributions at different speeds, respectively. In all cases, Bonferroni’s post-hoc t-test was used for multiple comparisons. Comparisons with p ≤ 0.05 were considered significant.

## Results

### Joint Angle Kinematics

Angular kinematics of different joints were similar in the left and right limbs (p>0.05, Figure 3a); therefore, parameters were compared across speed ranges using the data from the left hind limb only (Table 1). The left hind limb was chosen because the maximum and minimum joint angles occurred in the middle of the gait cycle, making the comparisons more reliable.

**Table 1.** Comparison of different parameters of overground walking at each speed range (Mean ± standard error).

| Measure | Parameter | 0.4-0.59<br>(m/s) | 0.6-0.79<br>(m/s) | 0.8-0.99<br>(m/s) | 1-1.19<br>(m/s) | 1.2-1.39<br>(m/s) | 1.4-1.6<br>(m/s) | Sig |
| --- | --- | --- | --- | --- | --- | --- | --- | --- |
| Hip joint<br>(n=6) | Minimum angle (deg) | 94.7±2.5<sup>a</sup> | 92.8±2.4<sup>ab</sup> | 92.0±2.8<sup>bc</sup> | 93.4±2.8<sup>a</sup> | 94.9±2.7<sup>a</sup> | 94.5±3.0<sup>ac</sup> | * |
| | Maximum angle (deg) | 109.8±4.1<sup>a</sup> | 111.4±1.0<sup>a</sup> | 111.8±3.8<sup>a</sup> | 112.0±3.7<sup>a</sup> | 111.9±3.8<sup>a</sup> | 112.2±3.7<sup>a</sup> | - |
| | ROM (deg) | 15.0±2.7<sup>a</sup> | 18.6±2.1<sup>a</sup> | 19.8±1.7<sup>a</sup> | 18.6±1.7<sup>a</sup> | 17.0±1.7<sup>a</sup> | 17.7±1.4<sup>a</sup> | - |
| | Gait percentage of max angle | 57.3±2.6<sup>a</sup> | 53.0±1.6<sup>a</sup> | 52.3±0.9<sup>a</sup> | 50.0±0.8<sup>a</sup> | 47.0±1.9<sup>a</sup> | 46.0±2.8<sup>a</sup> | - |
| | Gait percentage of min angle | 94.6±1.9<sup>a</sup> | 94.3±0.8<sup>a</sup> | 93.3±0.6<sup>a</sup> | 93.3±0.9<sup>a</sup> | 93.3±0.6<sup>a</sup> | 93.3±1.1<sup>a</sup> | - |
| Knee joint<br>(n=6) | Minimum angle (deg) | 107.6±2.5<sup>a</sup> | 105.1±2.6<sup>ab</sup> | 101.5±2.2<sup>bc</sup> | 99.5±2.4<sup>cd</sup> | 98.7±3.2<sup>bd</sup> | 99.2±2.9<sup>bd</sup> | ** |
| | Maximum angle (deg) | 137.7±3.8<sup>a</sup> | 144.5±2.2<sup>a</sup> | 145.5±2.4<sup>a</sup> | 146.3±2.6<sup>a</sup> | 146.0±2.7<sup>a</sup> | 148.3±3.3<sup>a</sup> | ** |
| | ROM (deg) | 30.1±6.5<sup>ac</sup> | 39.7±1.3<sup>abc</sup> | 43.9±0.9<sup>ac</sup> | 46.8±0.7<sup>b</sup> | 47.2±0.9<sup>abc</sup> | 49.1±0.7<sup>abc</sup> | ** |
| | Gait percentage of max angle | 98.3±0.8<sup>a</sup> | 94.0±0.8<sup>b</sup> | 91.6±0.3<sup>bc</sup> | 90.6±0.4<sup>abc</sup> | 89.3±0.4<sup>bc</sup> | 88.6±0.6<sup>c</sup> | ** |
| | Gait percentage of min angle | 73.3±0.6 | 68.3±0.9<sup>a</sup> | 65.6±0.9<sup>b</sup> | 64.3±0.9<sup>b</sup> | 63.6±1.4<sup>ab</sup> | 63.6±1.2<sup>ab</sup> | ** |
| Ankle joint<br>(n=6) | Minimum angle (deg) | 120.1±6.0<sup>a</sup> | 117.1±5.2<sup>a</sup> | 113.7±5.4<sup>b</sup> | 109.7±5.5<sup>c</sup> | 107.8±5.5<sup>bc</sup> | 107.7±5.3<sup>bc</sup> | ** |
| | Maximum angle (deg) | 145.2±3.5<sup>ab</sup> | 148.2±3.9<sup>ab</sup> | 150.1±3.6<sup>a</sup> | 151.3±3.4<sup>ab</sup> | 151.3±3.0<sup>ab</sup> | 152.2±3.5<sup>b</sup> | ** |
| | ROM (deg) | 25.1±3.1<sup>a</sup> | 31.0±2.5<sup>a</sup> | 36.4±2.7<sup>b</sup> | 41.5±2.7<sup>c</sup> | 43.4±3.6<sup>bc</sup> | 44.5±2.5<sup>c</sup> | ** |
| | Gait percentage of max angle | 55.6±0.9<sup>a</sup> | 52.3±1.2<sup>a</sup> | 49.6±1.3<sup>ab</sup> | 45.3±1.1<sup>bc</sup> | 42.3±1.2<sup>bc</sup> | 40.3±1.4<sup>bc</sup> | ** |
| | Gait percentage of min angle | 81.0±1.1 | 77.0±0.6<sup>a</sup> | 73.6±0.6<sup>ab</sup> | 71.3±0.6<sup>b</sup> | 69.6±0.6<sup>b</sup> | 69.0±0.8<sup>b</sup> | ** |
| MTP<br>(n=6) | Gait percentage of max height | 80.7±0.01<sup>a</sup> | 75.7±0.01<sup>b</sup> | 70.3±0.01<sup>c</sup> | 66.0±0.02<sup>bcd</sup> | 62.0±0.02<sup>cd</sup> | 58.7±0.01<sup>d</sup> | ** |
| | ROM (mm) | 43.8±2.8<sup>a</sup> | 46.5±2.1<sup>a</sup> | 46.3±3.1<sup>a</sup> | 48.5±3.2<sup>a</sup> | 53.2±4.1<sup>a</sup> | 56.1±3.5<sup>a</sup> | - |
| Toe<br>(n=3) | Gait percentage of max height | 89.3±0.02<sup>a</sup> | 87.3±0.04<sup>a</sup> | 73.3±0.9<sup>a</sup> | 71.3±0.08<sup>a</sup> | 68.0±0.1<sup>a</sup> | 66.0±0.1<sup>a</sup> | - |
| | ROM (mm) | 20.8±2.1<sup>a</sup> | 22.1±2.0<sup>a</sup> | 22.3±1.2<sup>a</sup> | 25.0±1.2<sup>a</sup> | 26.6±2.0<sup>a</sup> | 30.1±3.9<sup>a</sup> | - |
| Pelvis<br>(n=6) | ROM (mm) | 20.3±1.7<sup>a</sup> | 24.6±0.6<sup>a</sup> | 23.7±1.2<sup>a</sup> | 25.4±0.9<sup>ab</sup> | 28.8±0.9<sup>ab</sup> | 33.5±1.6<sup>b</sup> | * |
| | Hip-Hike ROM (mm) | 24.3±3.8<sup>ab</sup> | 31.2±2.7<sup>ab</sup> | 28.7±2.3<sup>ab</sup> | 21.2±1.8<sup>a</sup> | 14.3±1.6<sup>a</sup> | 17.7±2.3<sup>a</sup> | ** |
| Stride data | Stride length (m) | 0.52±0.10 | 0.58±0.005 | 0.63±0.007 | 0.66±0.007 | 0.70±0.006 | 0.74±0.008 | ** |
| | Swing time (s) | 0.36±0.003 | 0.33±0.001 | 0.30±0.001 | 0.28±0.0001 | 0.27±0.001<sup>a</sup> | 0.27±0.001<sup>a</sup> | *** |
| | Stance time (s) | 0.66±0.006 | 0.52±0.001 | 0.42±0.001 | 0.34±0.001 | 0.27±0.002 | 0.23±0.002 | *** |
| | Stride time (s) | 1.02±0.007 | 0.86±0.002 | 0.73±0.001 | 0.62±0.001 | 0.54±0.001 | 0.50±0.001 | *** |
| | Duty factor (four limbs) | 0.65±0.003 | 0.61±0.001 | 0.58±0.001 | 0.55±0.001 | 0.50±0.002 | 0.46±0.003 | *** |
|  | Duty factor |  |  |  |  |  |  |  |
| | Hindlimbs | 0.63±0.003 | 0.60±0.001 | 0.57±0.001 | 0.53±0.002 | 0.46±0.002 | 0.43±0.002 | *** |
| | forelimbs | 0.66±0.003 | 0.62±0.001 | 0.60±0.001 | 0.57±0.002 | 0.54±0.002 | 0.49±0.002 | *** |
| coupling | Diagonal coupling (%) | 24.2±0.33<sup>a</sup> | 23.6±0.11<sup>a</sup> | 22.0±0.10 | 18.6±0.19 | 12.4±0.34 | 7.2±0.38 | *** |
| | Homolateral coupling (%) | 75.3±0.35 | 73.7±0.09 | 71.6±0.10 | 67.6±0.19 | 60.4±0.31 | 54.3±0.27 | *** |
| | Homologous coupling (%) | 49.4±0.41<sup>a</sup> | 50.2±0.12<sup>a</sup> | 50.2±0.09<sup>a</sup> | 50.4±0.09<sup>a</sup> | 50.2±0.14<sup>a</sup> | 50.2±0.14<sup>a</sup> | - |
| Cyclogram Area | Left knee vs ankle (deg<sup>2</sup>) | 562.45±61.6<sup>a</sup> | 691.56±55.5<sup>abcd</sup> | 849.49±48.9<sup>ef</sup> | 911.06±73.9<sup>be</sup> | 849.31±82.1<sup>cf</sup> | 783.89±63.7<sup>df</sup> | * |
| | Left hip vs ankle (deg<sup>2</sup>) | 206.61±57.5<sup>a</sup> | 235.29±64.9<sup>a</sup> | 290.69±74.5<sup>a</sup> | 294.47±73.1<sup>a</sup> | 259.06±61.7<sup>a</sup> | 241.05±55.4<sup>a</sup> | - |
| | Left hip vs knee (deg<sup>2</sup>) | 187.27±48.4<sup>a</sup> | 213.50±52.8<sup>a</sup> | 250.28±61.1<sup>a</sup> | 254.59±63.2<sup>a</sup> | 230.54±55.7<sup>a</sup> | 227.28±54.6<sup>a</sup> | - |
| | Right knee vs ankle (deg<sup>2</sup>) | 680.18±43.0<sup>a</sup> | 758.31±42.1<sup>ac</sup> | 899.30±32.4<sup>c</sup> | 966.31±59.3<sup>c</sup> | 909.96±45.2<sup>c</sup> | 889.01±54.3<sup>ac</sup> | * |
| | Right hip vs ankle (deg<sup>2</sup>) | 225.91±33.5<sup>a</sup> | 252.97±46.7<sup>a</sup> | 290.89±58.2<sup>a</sup> | 282.43±48.6<sup>a</sup> | 264.46±31.2<sup>a</sup> | 252.49±28.6<sup>a</sup> | - |
| | Right hip vs knee (deg<sup>2</sup>) | 190.05±36.8<sup>a</sup> | 212.86±51.8<sup>a</sup> | 232.62±61.4<sup>a</sup> | 231.87±54.0<sup>a</sup> | 215.91±47.2<sup>a</sup> | 205.16±45.9<sup>a</sup> | - |
| Cyclogram regularity | Left knee vs ankle | 0.40±0.040<sup>a</sup> | 0.50±0.026<sup>ab</sup> | 0.53±0.023<sup>b</sup> | 0.53±0.020<sup>ab</sup> | 0.51±0.028<sup>ab</sup> | 0.53±0.026<sup>ab</sup> | * |
| | Left hip vs ankle | 0.35±0.044<sup>a</sup> | 0.45±0.022<sup>ac</sup> | 0.49±0.026<sup>c</sup> | 0.50±0.017<sup>ac</sup> | 0.49±0.022<sup>ac</sup> | 0.52±0.027<sup>ac</sup> | * |
| | Left hip vs knee | 0.38±0.034<sup>a</sup> | 0.47±0.028<sup>ab</sup> | 0.51±0.025<sup>b</sup> | 0.52±0.018<sup>ab</sup> | 0.52±0.023<sup>ab</sup> | 0.55±0.032<sup>ab</sup> | * |
| | Right knee vs ankle | 0.33±0.042<sup>a</sup> | 0.48±0.032<sup>b</sup> | 0.52±0.022<sup>b</sup> | 0.53±0.019<sup>ab</sup> | 0.52±0.018<sup>ab</sup> | 0.53±0.023<sup>ab</sup> | * |
| | Right hip vs ankle | 0.28±0.035<sup>a</sup> | 0.41±0.034<sup>b</sup> | 0.45±0.026<sup>b</sup> | 0.49±0.016<sup>b</sup> | 0.49±0.015<sup>ab</sup> | 0.51±0.019<sup>b</sup> | * |
| | Right hip vs knee | 0.34±0.044<sup>a</sup> | 0.48±0.029<sup>ab</sup> | 0.51±0.023<sup>b</sup> | 0.53±0.015<sup>ab</sup> | 0.52±0.017<sup>ab</sup> | 0.53±0.022<sup>ab</sup> | * |

a,b,c,d mean values not sharing superscripts differ significantly with Bonferroni adjustment
\*p < 0.05, \*\*p < 0.001, \*\*\*p < 0.0005

**Figure 3.**
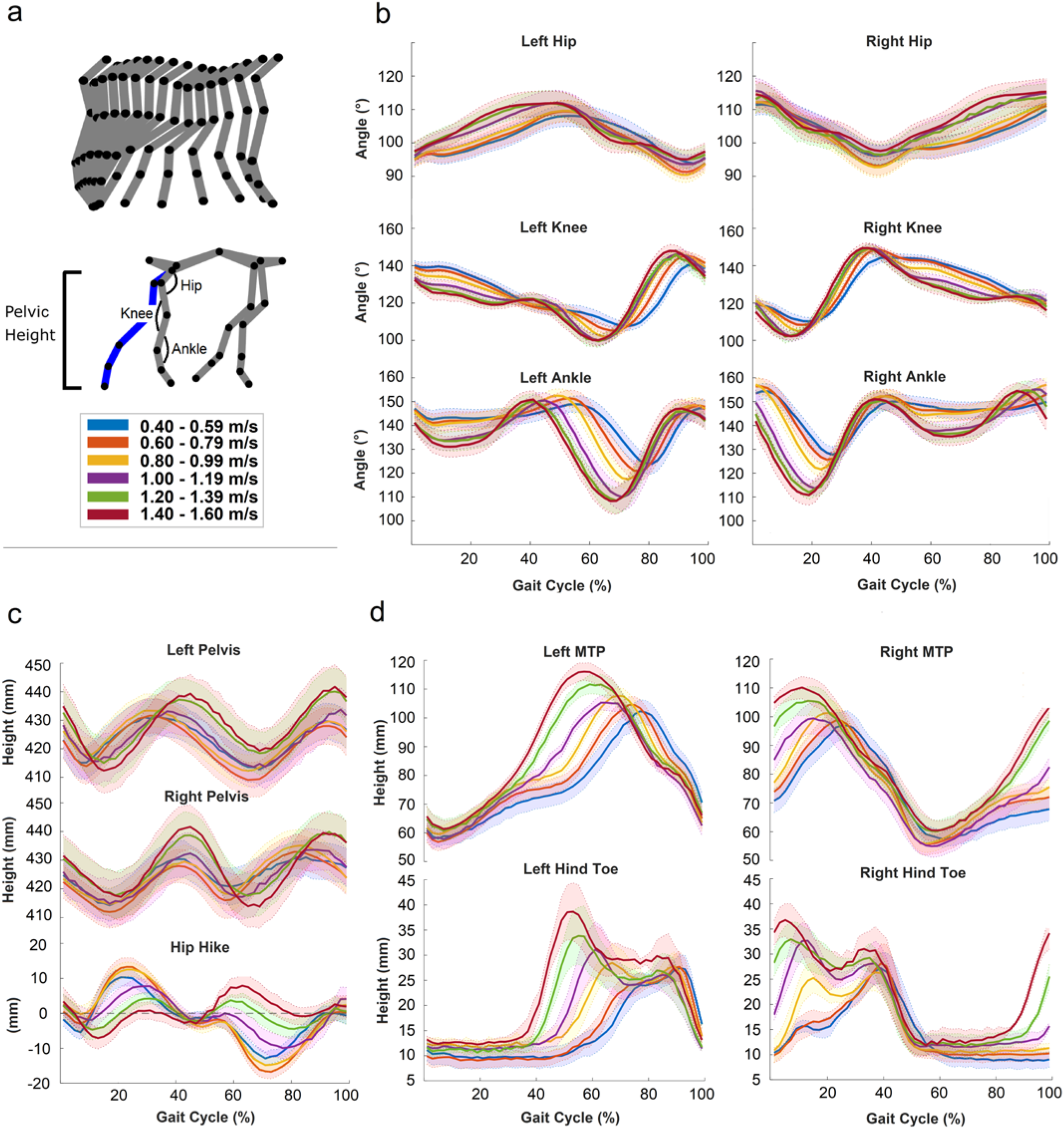
Summary of hindlimb kinematics for overground walking at different speeds. **(a)** Left hindlimb stick figures showing the changes in joint angles during the gait cycle and how the angles were calculated across six different speed ranges. **(b)** Mean joint angles of hip, knee, and ankle of left (left column) and right (right column) limbs (n= {9, 9, 9, 6, 6, 6} – n represents the number of animals included in the average joint angles at 0.4-0.6 m/s, 0.6-0.8 m/s, 0.8-1.0 m/s, 1.0-1.2 m/s, 1.2-1.4 m/s, and 1.4-1.6 m/s, respectively). **(c)** Change in the left and right pelvic heights and hip-hike (left pelvic height – right pelvic height) with speed (n= {9, 9, 9, 6, 6, 6}). **(d)** Clearance over ground of the MTP joint n= {9, 9, 9, 7, 6, 6} and hind toe n= {4, 4, 4, 3, 3, 3} for the left and right limbs

For the ankle and knee joints, as the speed of walking increased, the crests and troughs in the gait cycle increased and decreased, respectively (p<0.001, Table 1). Consequently, the range of motion (ROM) for the knee joint increased from 30.1±6.5° at the 0.4-0.59m/s speed to 39.7±1.3°, 43.9±0.9°, 46.8±0.7°, 47.2±0.9°, and 49.1±0.7° (mean±SE) at the 0.6-0.79m/s, 0.8-0.99m/s, 1.0-1.19m/s, 1.2-1.39m/s, and 1.4-1.6m/s speeds, respectively (p<0.001). The ankle joint ROM increased from 25.1±3.1° at the 0.4-0.59m/s speed to 31.0±2.5°, 36.4±2.7°, 41.5±2.7°, 43.4±3.6°, and 44.5±2.5° at the 0.6-0.79m/s, 0.8-0.99m/s, 1.0-1.19m/s, 1.2-1.39m/s, and 1.4-1.6m/s speeds, respectively (p<0.001). In addition, the time points within the gait cycle at which maximal extension and maximal flexion occurred differed significantly at different speeds for both the knee and ankle, occurring earlier in the gait cycle as the speed increased (knee max: p<0.001; knee min: p<0.001; ankle max: p<0.001; ankle min: p<0.001). For the knee joint, the percent of the gait cycle at which the crests and troughs occurred moved from 98.3±0.8% (mean±SE) to 88.6±0.6% and from 73.3±0.6% to 63.6±1.2% with increasing speed from 0.4-0.59m/s to 1.4-1.6m/s, respectively (Figure 3a and Table 1). Similarly, for the ankle joint, these points moved from 55.6±09% (mean±SE) to 40.3±1.4% and from 81.0±1% to 69.0±0.8%. For the hip joint, only the percent of gait cycle at which the maximal flexion occurred was significantly different with speed (p<0.001).

### Vertical Displacement of the Pelvis and Toe

The vertical displacement of the left and right pelvic markers above the ground and hip-hike are shown in Figure 3c. Hip-hike was calculated by subtracting the left and right pelvic displacement at each point in the gait cycle. Left pelvic ROM and range of hip-hike were significantly different at different speed ranges (Table 1). The pelvic ROM increased from 20.3±1.7 mm to 33.5±1.6 mm with increasing speed from 0.4-0.59m/s to 1.4-1.6m/s (p<0.001). In addition, hip-hike ROM decreased from 24.3±3.8 for the 0.4-0.6m/s speed to 17.7± 2.3 for the 1.4-1.6m/s (p<0.001).

The moment at which the MTP reached its maximal height over the ground occurred significantly earlier (p<0.001) at higher speeds. The maximal height of the MTP joint occurred at 80.7±0.01% of the gait cycle at the lowest speed range; this point shifted to 58.7±0.01% at the highest speed range. The vertical ROM of MTP joint and left hind-toe as well as the percent of the gait cycle where maximal clearance above the ground occurred for the hind-toe were similar across speeds.

### Gait Cycle Parameters

The mean stride length calculated across all animals showed statistically significant differences at all different speed ranges (p<0.001, Table 1, Figure 4a). The mean stride length increased from 0.52±0.10m for the 0.4-0.59m/s speed to 0.58±0.005m, 0.63±007m, 0.66±0.007m, 0.70±0.006m, and 0.74±0.008m (mean±SE) for the 0.6-0.79m/s, 0.8-0.99m/s, 1.0-1.19m/s, 1.2- 1.39m/s, and 1.4-1.6m/s speeds, respectively. No significant variation between the four limbs was observed at any of the speed ranges.

**Figure 4.**
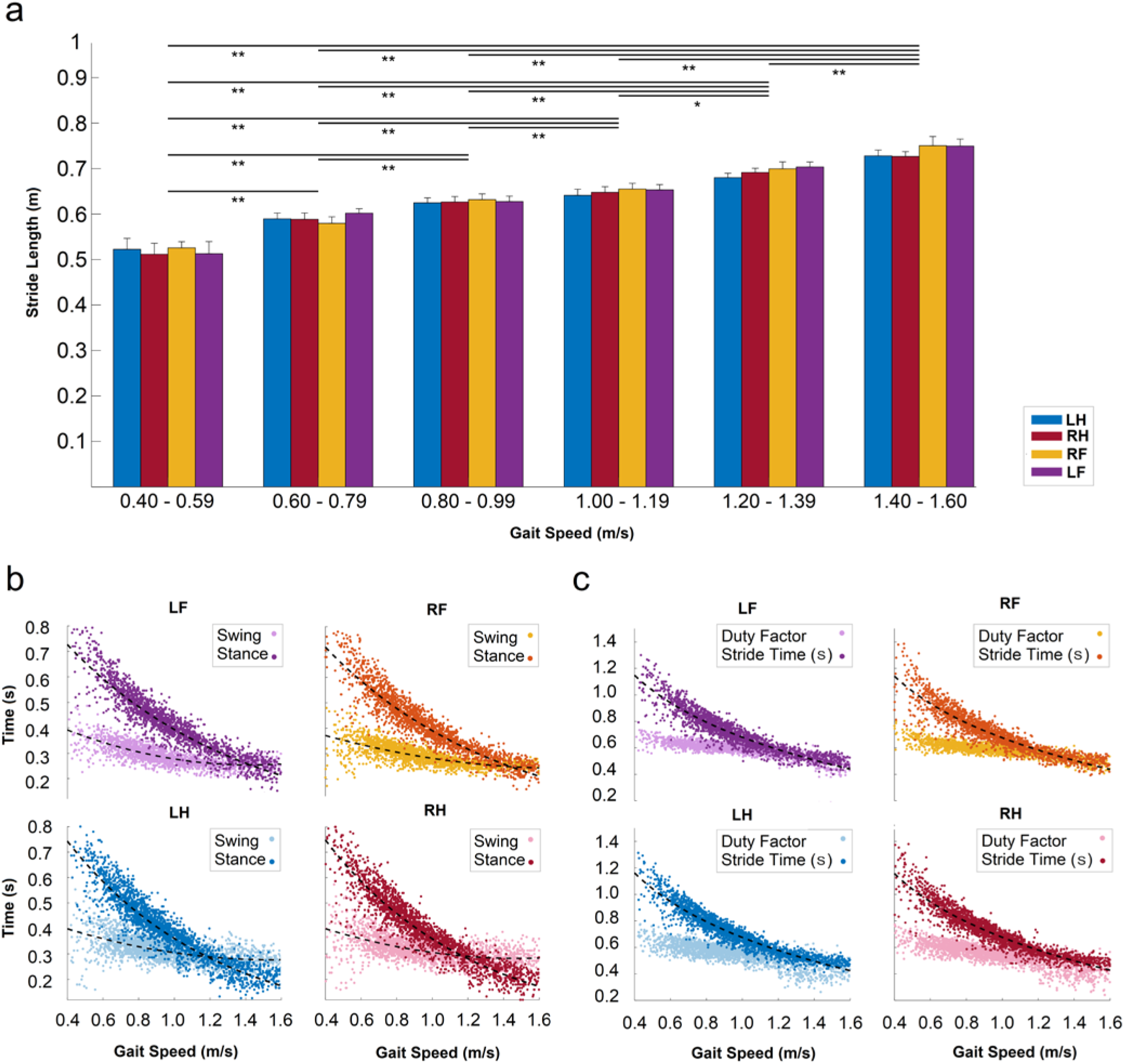
Summary of gait parameters for different overground walking speeds. **(a)** Mean values of stride length for all four limbs at different speed ranges (mean ± standard error; * p<0.05, **p<0.001, with Bonferroni correction). **(b)** The duration of swing time and stance time for all four limbs with curves fitted (fit equations for swing phases, LF: y= 0.132 x2 - 0.375 x + 0.521 (r = 0.53), RF: y= 0.072 x^2^ - 0.250 x + 0.460 (r = 0.65), LH: y= 0.092 x^2^ - 0.285 x + 0.498 (r =0.56), RH: y= 0.099 x^2^ - 0.294 x + 0.501 (r = 0.52) and stance phase, LF: y= 1.095 e^(−1.023 x)^ (r = 90), RF: y= 1.081 e^(−1.019 x)^ (r = 0.90), LH: y= 1.203 e^(−1.204 x)^ (r = 0.91), RH: y= 1.221 e^(−1.223 x)^ (r = 0.91)). **(c)** The duration of stride time and duty factor values for all four limbs with curves fitted for the stride time (LF: y= −0.511 ln(x)+0.680 (r = 0.91), RF: y= −0.508 ln(x)+0.676 (r = 0.91), LH: y= −0.532 ln(x)+0.675 (r = 0.93), RH: y= −0.526 ln(x)+0.676 (r = 0.92)). LF – left forelimb; RF – right forelimb; LH – left hindlimb; RH – right hindlimb.

Both the swing and stance times decreased significantly (p<0.0005) with increasing speed. The mean swing time changed from 0.36±0.003s to 0.33±0.001s, 0.30±0.001s, 0.28±0.0001s, 0.27±0.001s, and 0.27±0.001s with increasing speed ranges and the mean stance time for all animals decreased from 0.66±0.006s to 0.52±0.001s, 0.42±0.001s, 0.34±0.001s, 0.27±0.002s, and 0.23±0.002s. The duty factor decreased significantly (p<0.0005) with increasing speed from 0.65±0.003, to 0.61±0.001, 0.58±0.001, 0.55±0.001, 0.50±0.002, and 0.46±0.003. Furthermore, the averaged duty factor showed significant differences (p<0.001) between forelimbs (0.54 ± 0.001) and hindlimbs (0.58 ± 0.001).

The mean stride time significantly decreased with increasing speed (p<0.0005). A decrease from 1.02±0.007s to 0.86±0.002s, 0.73±0.001s, 0.62±0.001s, 0.54±0.001s, and 0.50±0.001s was observed for mean stride time with increasing speed; the differences were significant between all possible pairs at various speed ranges (Figure 4c and Table 1)

### Interlimb and Inter-joint Coordination

Interlimb coupling was calculated between the four limbs (Figure 5a). While diagonal (Figure 5b) and homolateral (Figure 5d) coupling showed significant differences between the various speed ranges (p<0.0005), no significant differences for homologous coupling (Figure 5c) was detected between speed ranges. Diagonal coupling decreased with increasing speed from 24.2±0.33% to 23.6±0.11%, 22.0±0.10%, 18.6±0.19%, 12.4±0.34%, and 7.2±0.38%. Similarly, homolateral coupling decreased from 75.3±0.35% to 73.7±0.09%, 71.6±0.10%, 67.6±0.19%, 60.4±0.31% and 54.3±0.27% with increasing walking speed.

**Figure 5.**
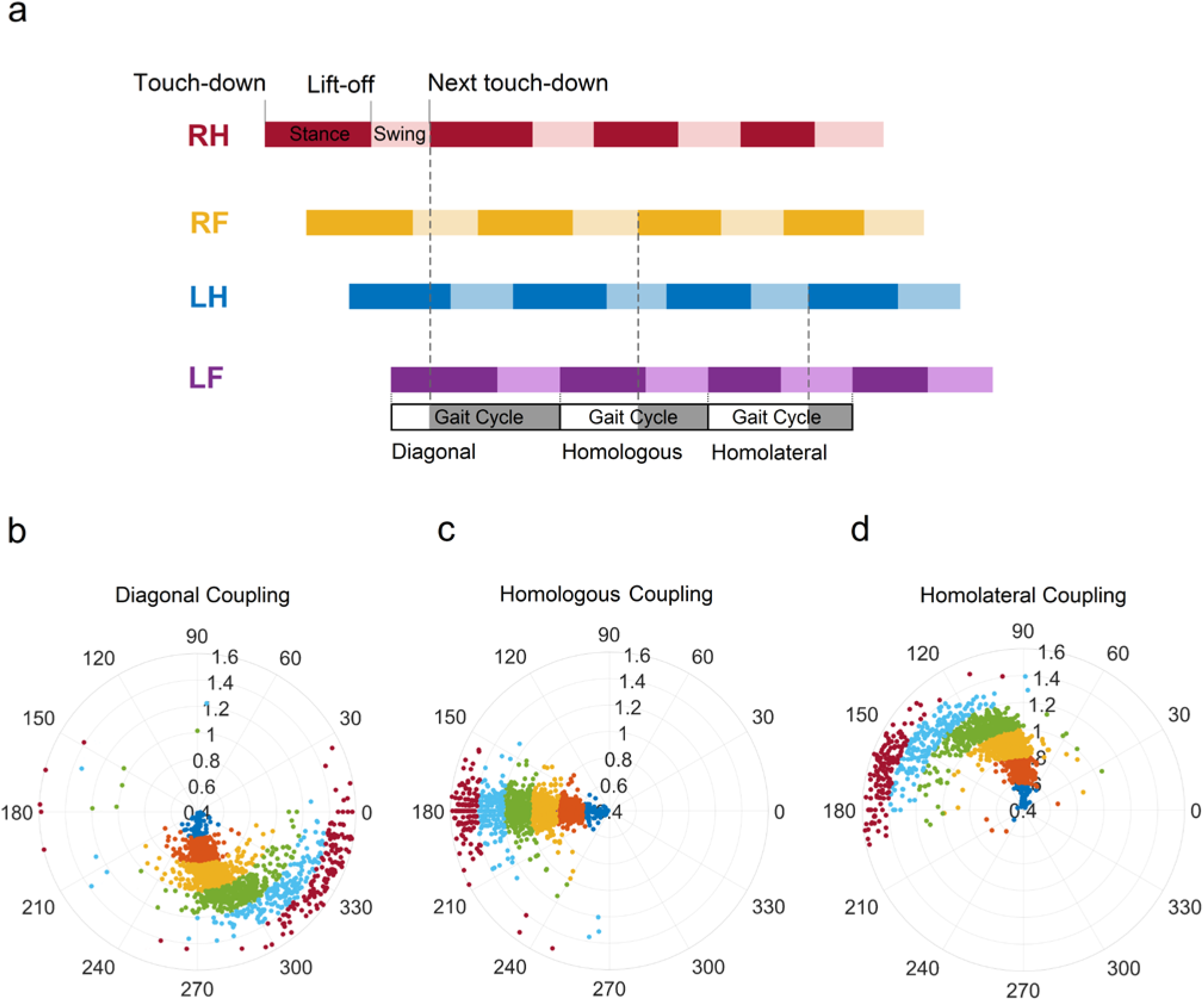
Summary of Interlimb coupling during overground walking at different speeds. **(a)** Duty cycle diagram shows how the coupling between the different limbs were calculated (stepping sequences are plotted for the 0.69 m/s overground walking speed). **(b)** Diagonal coupling between the right hindlimb (RH) and left forelimb (LF). At lower speeds, the RH leads the LF by 270°; this phase angle approaches 0° at higher speeds. **(c)** Homologous coupling of the right forelimb (RF) and LF. The RF consistently leads the LF by 180° at all speeds. **(d)** Homolateral coupling between the left hindlimb (LH) and LF. The LH leads the LF by 90°, although this phase angle approaches 180° at higher speeds. Gait transitions from a lateral sequence to a diagonal sequence at walking speed of around 1.0 m/s (b and d).

Cyclograms of knee-ankle, hip-ankle, and hip-knee inter-joint coordination during the gait cycle are shown for one representative pig (Figure 6a) and averaged across all pigs (Figure 6b). Table 1 shows the mean area of the cyclograms of various joints at different speed ranges. The left knee vs ankle cyclograms mean area increased from 562.45±61.6(deg^2^) to 691.56±55.5(deg^2^), 849.49±48.9(deg^2^), and 911.06±73.9(deg^2^) with increasing speed range from 0.4-0.59m/s to 1-1.19m/s and decreased to 849.31±82.1(deg^2^) and 783.89±63.7(deg^2^) at speed ranges of 1.2-1.39m/s and 1.4-1.59m/s, respectively (p<0.05). Similarly, the mean area of the right knee vs ankle increased from 680.18±43.0(deg^2^) to 758.31±42.1(deg^2^), 899.30±32.4(deg^2^), and 966.31±59.3(deg^2^), as the overground walking speed increased from 0.4-0.59m/s to 1-1.19m/s and then decreased to 909.96±45.2(deg^2^) and 889.01±54.3(deg^2^) at the speed ranges of 1.2-1.39m/s and 1.4-1.59m/s, respectively (p<0.05).

**Figure 6.**
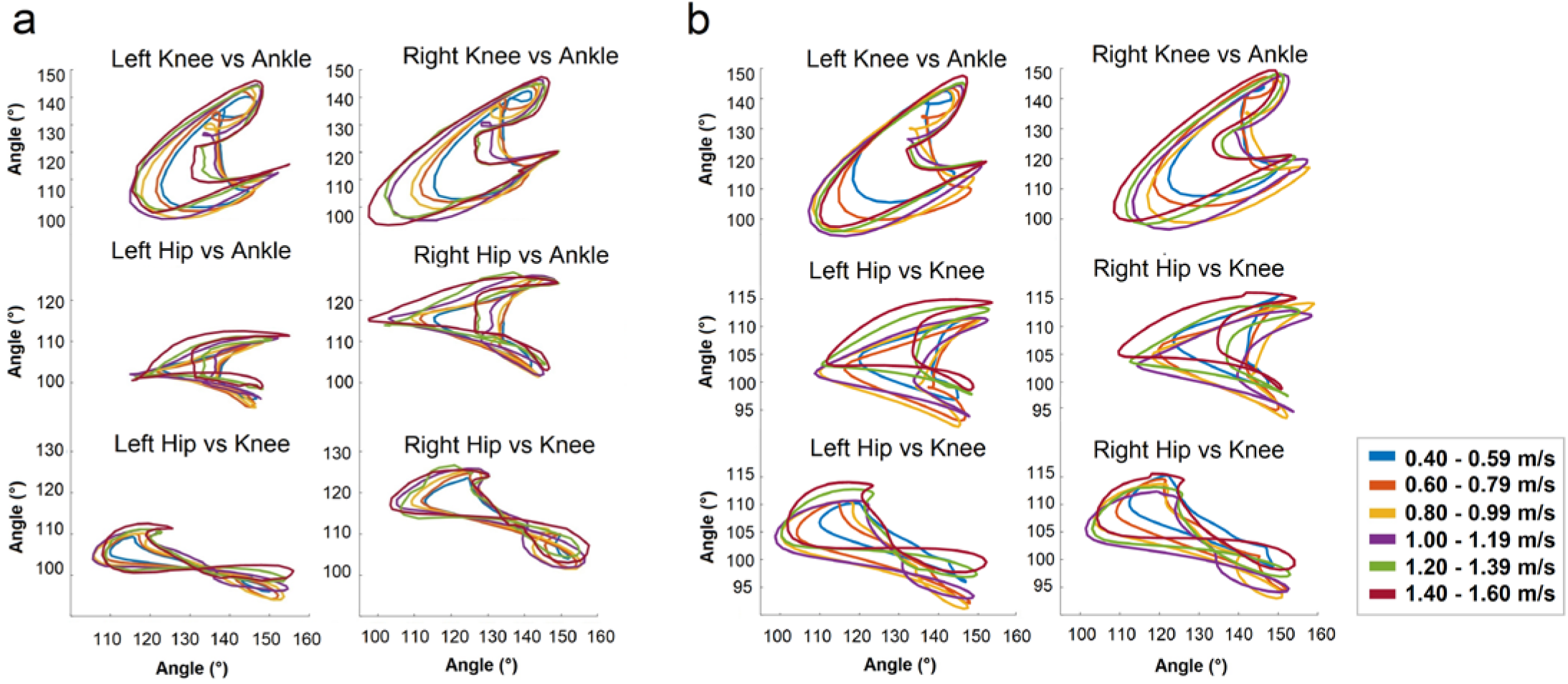
Cyclograms indicating inter-joint coupling in a representative animal **(a),** and average across all animals **(b)** for knee-ankle, hip-ankle, and hip-knee joints at each speed range.

Pairwise comparisons only demonstrated significant differences at given speeds for the area of left and right knee vs ankle cyclograms (p<0.05 with Bonferroni adjustment). The cyclogram areas of left knee vs ankle showed significant differences between 0.4-0.59m/s and 0.8-0.99m/s, 0.4-0.59m/s and 1.0-1.19m/s, 0.4-0.59m/s and 1.2-1.39m/s, 0.4-0.59m/s and 1.4-1.59m/s, 0.6-0.79m/s and 0.8-0.99m/s, 1.0-1.19m/s and 1.2-1.39m/s, and 1.0-1.19m/s and 1.4-1.59m/s. The right knee vs ankle cyclogram area showed significant differences between 0.4-0.59m/s and 0.8-0.99m/s, 0.4-0.59m/s and 1.0-1.19m/s, and 0.4-0.59m/s and 1.2-1.39m/s (Table 1).

Left knee vs hip cyclogram regularity values increased from 0.40±0.040 to 0.50±0.026, 0.53±0.023, 0.53±0.020, 0.51±0.028, and 0.53±0.026 with increasing speed from 0.4-0.59m/s to 1.2-1.4m/s (p<0.05). The left hip vs ankle regularity scores increased from 0.35±0.044 to 0.45±0.022, 0.49±0.026, 0.50±0.017, 0.49±0.022, and 0.52±0.027 with increasing speed of walking (p<0.05). In addition, the left hip vs knee cyclogram regularities increased with increasing walking speed from 0.38±0.034, to 0.47±0.028, 0.51±0.025, 0.52±0.018, 0.52±0.023, and 0.55±0.032. Furthermore, the right knee vs ankle, right hip vs ankle, and right hip vs knee cyclogram regularity values experienced similar increases as the walking speed increased from 0.4-0.59m/s to 1.2-1.4m/s (Table 1).

The regularities of cyclograms showed significantly different values across all speed ranges. The left knee vs ankle regularity values were significantly different between 0.4-0.59m/s and 0.8-0.99m/s speed ranges. For the right knee vs ankle cyclogram, the regularities were significantly different at 0.4-0.59m/s and 0.6-0.79m/s, and 0.4-0.59m/s and 0.8-0.99m/s speed ranges. For the hip vs ankle cyclograms, the regularities of the left side were significantly different between 0.4-0.59m/s and 0.8-0.99m/s, and for the right side, the regularity scores at 0.4-0.59m/s was significantly different from all other speed ranges except 1.2-1.39m/s. Lastly, both the right and left hip vs knee cyclograms had significantly different regularity scores between 0.4-0.59m/s and 0.8-0.99m/s speed ranges (Table 1).

### EMG Activity

An example of EMG activity recorded from the GM, BF and VL muscles in one pig is shown in Figure 7. Changes in the activation patterns, particularly the onset and offset points of EMG activity, were determined and compared across speed ranges. The low-activity regions shown with dashed green lines occurred 60-90% of the gait cycle for the left GM, 10-40% of the gait cycle for the right GM, 50-90% of the gait cycle for the left VL, 0-40% of the gait cycle for the right VL, 30-50% of the gait cycle for the left BF, and 80-100% of the gait cycle for the right BF.

**Figure 7.**
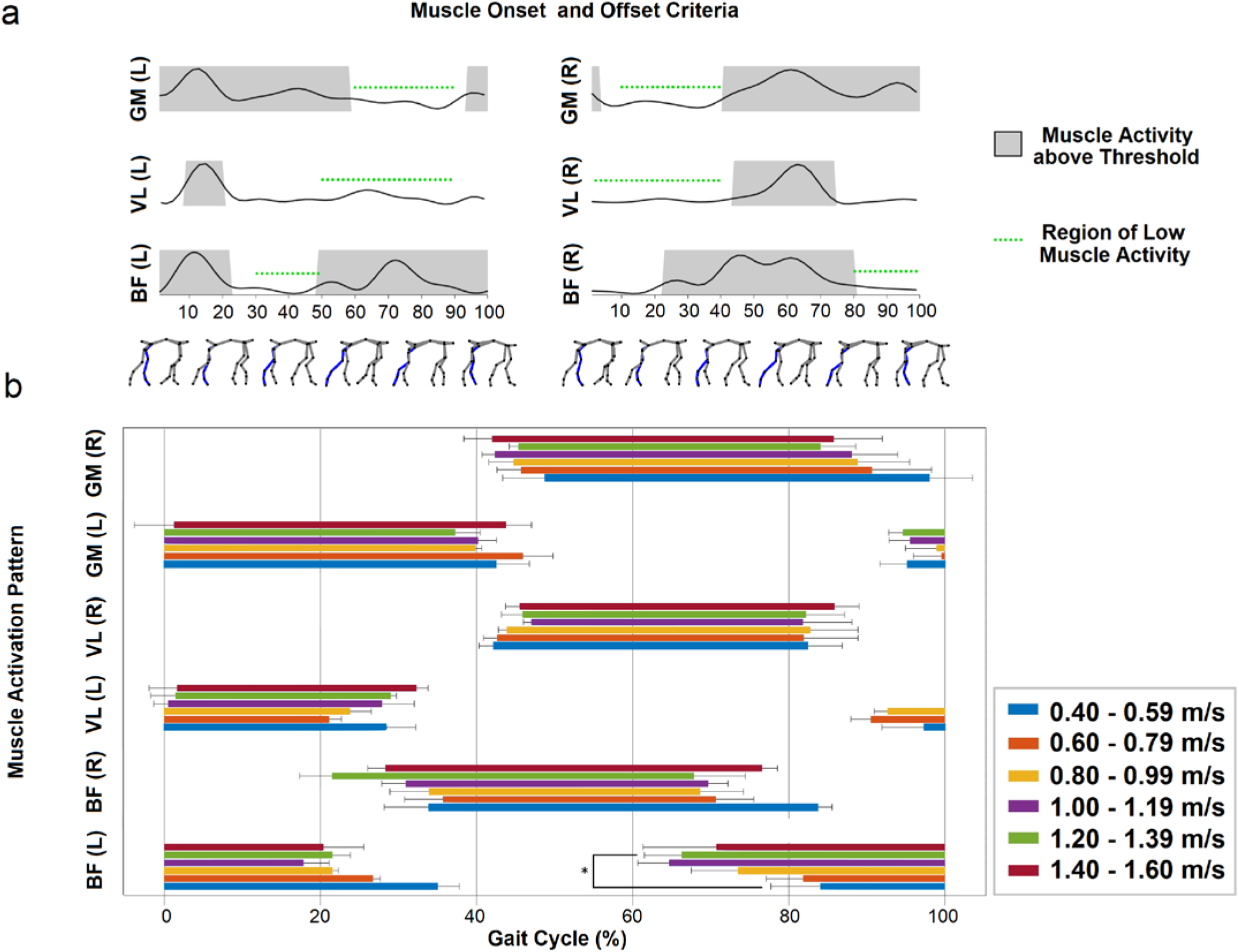
Summary of EMG activity during different speeds of overground walking. **(a)** Muscle activity of left and right gluteus medius (GM), biceps femoris (BF), and vastus lateralis (VL) muscles from one pig walking at 1.00-1.19 m/s. A muscle was considered to be active if the EMG activity exceeded a threshold identified based on low-activity region (represented by the green horizontal lines). **(b)** Average muscle activation patterns across 4 pigs, including onset and offset points, for each muscle across the gait cycle within each speed range.

Generally, all muscles exhibited one burst of activity during the gait cycle. The GM and VL muscles were activated primarily during the stance phase. For the BF muscles, activity began during the swing phase and terminated shortly after the beginning of the stance phase. The left BF muscle often exhibited a distinct two-burst pattern (Figure 7a), with the swing and stance bursts separated by a brief period of inactivity. In the right BF muscles, these two bursts were almost always overlapping and formed one sustained burst of activity. This left-right variability of the BF muscles may have been due to variable placement of the sEMG electrodes on each pig.

Figure 7b shows how muscle onset and offset points varied for strides within each speed range. At mid-speed (0.8-.0.99m/s), the mean onset and offset points (mean±SE) of each muscle were: 99±4% and 40±1% for the left GM, 45±3% and 89±7% for the right GM, 93±2% and 24±3% for the left VL, 44±1% and 83±6% for the right VL, 74±6% and 22±1% for the left BF, and 34±5% and 69±6% for the right BF. At different speeds, these values remained largely unchanged. Significant differences were only detected for the left BF onset and left BF offset points across the 6 speed ranges (p<0.05). Pairwise comparisons for the left BF onset showed significant differences at 0.4-0.6m/s (84±6%) and at 1.2-1.4m/s, (66±5%). Pairwise comparisons for the left BF offset values resulted in no significant differences.

## Discussion

This is the first study that focuses on analyzing the spatiotemporal parameters, kinematics, and EMG activity of overground walking in neurologically-intact adult YMPs. The results are critical for evaluating the role of sensorimotor neural circuits in controlling locomotion, and for quantifying the effects of neural injury and novel interventions on overground walking. The 6 different speed ranges chosen for gait analysis in this study encompass most of the speed ranges used in previous animal studies[22], [30]–[32]. Monitoring gait parameters over a wide range of overground walking speeds is particularly valuable not only for understanding the mechanisms of neural control during typical walking, but for understanding the manner in which locomotor patterns, kinematics, and muscle activation patterns change as a result of increasing walking speed.

### Overground Walking Parameters

A decrease in duration of step cycle was observed as the walking speed increased. Also, a significant decrease in swing time (~25% from 0.36±0.003s to 0.27±0.001s) was detected during increasing speed from 0.4-0.59m/s to 1.2-1.39m/s. Several previous studies in humans and animals have reported a similar effect of speed on the duration of the step cycle[22], [24], [31]– [35]. In addition, the effects of walking speed on swing time in YMPs obtained here are consistent with previous studies in cats[33], [36], [37], where a 21-22% reduction in swing time is reported as the cats speed up from walking to trotting.

The lateral sequence of footfall pattern (i.e., the limb to move after a hindlimb is the ipsilateral forelimb: RH, RF, LH, LF) shown in Figure 5a is also seen in gait patterns of other mammals such as cats, dogs, some primates, and infant/adult humans during crawling[34], [38]–[41]. Interestingly, a shift to a diagonal sequence was observed (RH, LF, LH, RF) at higher speeds in which the hindlimb was followed by the contralateral forelimb (Figure 5b). Generally, gait parameters showed higher variability in the lowermost walking speed ranges. A higher variability was also seen at the highest speed range; however, that was likely due to the lower number of strides taken at this speed. In addition, interlimb coordination indicated a gait transition at ~1m/s, confirming observations seen in treadmill-based walking for YMPs[22]. Thus, it is recommended that the speed range 0.8-1.19m/s be included when analyzing overground gait in YMPs.

Compared with treadmill-based walking of YMPs[22], the stride length for overground walking, measured at similar speeds to treadmill walking, was longer. However, a higher duty factor was reported in treadmill-based walking than in the overground walking in this study. Similar differences between overground walking and treadmill walking have been previously reported in humans and cats[24], [26], [42], [43]. In treadmill-based walking, once the treadmill belt starts moving, limbs are pulled back causing extension/flexion in different joints as well as tension in different muscles. These changes produce regular frequencies of afferent impulses from Golgi tendon organs, muscle spindles, and cutaneous receptors into the spinal central pattern generator (CPG), a network of neurons which controls locomotion. Furthermore, while the placement of a food tray in front of the pig during treadmill walking might influence head position, natural deviations in head position during overground walking often occur, inducing differences in neck and vestibular-ocular reflexes from those induced during treadmill walking. Finally, in contrast to treadmill walking, the pig does not need to adjust its walking velocity to match external measures (i.e., speed of treadmill belt) during overground walking. Nonetheless, in overground walking, several different descending inputs such as visual and vestibular signals influence the CPG, causing more irregular walking patterns[26].

### Interlimb and Inter-joint Coordination

Interlimb coordination is an important component for maintaining dynamic stability of movement and to satisfy the demands of various tasks (such as gait pattern transitions) in constantly changing environments[44]. It is well established that limb coordination during locomotion is governed by the CPG[35], [45]. Studies in mammalian quadrupeds demonstrated that movement of each of the four limbs is coordinated by separate spinal neuronal networks[46]. The main elements of forelimb and hindlimb CPGs are located at low cervical-high thoracic and upper to mid-lumbar spinal segments, respectively[47]. In addition, a distribution of propriospinal pathways communicate between the CPGs controlling fore/hindlimbs through long/short and ascending/descending pathways[48]. Somatosensory feedback has an important role in the activation of inhibitory and excitatory propriospinal tracts[49]. These pathways which project diagonally and homolateraly to the neural networks, influence the locomotor pattern by modulating forelimb/hindlimb coordination, gait transition, and adjustments in trajectory in response to perturbations[50], [51]. Supraspinal descending signals also control limb coordination and the temporal sequence of muscle activity in the four limbs. Studies in YMPs have shown that stimulation of the mesencephalic locomotor region can evoke responses in extensor and flexor muscles of all limbs and generate various gait patterns and speeds[15].

In the current study, a gait transition from walking to trotting occurred at speeds around 1.0m/s. However, the 180 degree phase difference between left and right limbs remained unchanged, similarly to what has been reported in overground locomotion of cats[52]. The homologous coupling showed no further gait transition (from symmetrical gait patterns such as walk or trot to asymmetrical gaits like pace, gallop, or bound) at the speeds tested. These results can be used for evaluating the functionality of neural and biomechanical mechanisms ruling interlimb coordination in injured YMPs.

Cyclograms are useful for diagnosis and tracking recovery following surgery or SCI[22], [53]. They have been used to qualitatively analyze inter-joint coordination in animal studies[30]. The cyclogram shapes are highly responsive to walking speed and abnormalities[54]. Recently, Boakye et al.[22] reported no difference in joint angles (distal and proximal), and no differences in cyclogram patterns, across varying speeds for treadmill-based walking of YMPs; this indicated that the inter-joint coordination was consistent from step to step. However, the cyclograms for treadmill walking of Japanese macaque monkeys showed that while the temporal components of locomotion were well-coordinated, there were differences in the pattern of the gait[55]. In the present study, as a general trend, the areas associated with all cyclograms increased from the slowest speed 0.4-0.59m/s to the 0.8-0.99m/s speed range. Generally, as the speed of walking increases the joint ROMs increases, leading to an increase in the area of the cyclograms[54], which was also seen in this study. This has also been shown in humans where the area of the hip vs knee cyclograms increases linearly with the walking speed[56].

Regularity measures provide a quantitative assessment of the relationship between step cycles and provide information on the mechanisms underlying the control of movement[29]. Regularity of cyclograms has been used to examine changes in gait following rehabilitation programs in persons with incomplete SCI; the higher post-training regularity scores indicated higher consistency in the walking pattern [57]. Cyclogram regularity has been also used as a measure for comparing the walking patterns of neurologically-intact study participants and participants with incomplete SCI[58]. Neurologically-intact participants had higher regularity scores compared with participants with incomplete SCI. In addition, the hip-knee cyclogram regularity values increased in both groups when moving from slow to faster walking speeds[58].

### Angular Joint Kinematics

Four subphases defined for a gait cycle, F and E1 (swing phase), and E2 and E3 (stance phase) were originally proposed by Philippson[59]. At the hip joint, these four subphases can be divided into two major phases of extension and flexion[37] (Figure 3b). At the onset of the stance phase (0% gait cycle for the left hindlimb), the ankle flexes a few degrees to preserve limb progression, the knee flexes to absorb the shock in loading response to the weight acceptance, and the hip joint extends (E2). During midstance, the ankle and knee joints extend (E3); for the ankle and hip joints the maximum extension occurs ~40%-60% of the gait cycle. At the onset of the swing phase (F) flexion of the hip, knee, and ankle takes place to elevate the limb for toe-off (Figure 3b, 1d) and advance the limb through space. The flexion of joints continues until the middle of the swing phase (F). In the final advancement of the limb forward (E1) the knee and ankle joints extend, the limb is lowered for another touchdown to start the next step cycle. In the present study, knee flexion at the start of the swing phase (F) showed a faster rate of flexion at higher speeds of walking (steeper slope can be seen for left knee joint angles in Figure 3b), while in the E1 phase the knee and ankle extension at different speeds of walking occurred at the same rate.

Hip-hike curves show that at speed ranges from 0.4-0.59m/s to 0.8-0.99m/s the difference between left or right pelvic vertical displacement increases for higher speeds. At speeds greater than 1.0m/s however, the absolute values of the hip-hike begin to decrease. These findings further support the transition in gait patterns at ~1.0m/s. During symmetrical (walk, trot or pace) to asymmetrical (bound or gallop) gait transition, the pelvic girdles begin to move synchronously[44]. Although here we did not detect any asymmetrical gait patterns, the left and right pelvises showed in-phase vertical movement at speeds greater than 1.0m/s.

### Muscle Activation Pattern

The GM and VL muscles primarily exhibited a single burst of activity during the gait cycle at the beginning of the stance phase. Although no significant differences were detected in the EMG activity onset or offset points for the left and right GM and VL muscles, or right BF muscle, the left BF muscle did experience a significantly earlier onset point at faster walking speeds (1.2-1.39m/s) when compared to walking at slower speeds (0.4-0.59m/s). Considerable EMG variability was observed across all pigs, likely due to variable placement of the sEMG electrodes from animal to animal. The activity pattern of GM and VL aligned well with activity seen in many species, where different muscles contribute synergistically to flexion or extension around different joints[12], [45], [60], [61]. These similarities likely owe to conserved evolutionary development of the CPG circuits which drive locomotion in vertebrates[62]–[65]. Others have noted that at higher velocities, muscle activation peaks tend to undergo a phase shift similar to that of the toe-off time, occurring earlier in the gait cycle[60] [60]. Lastly, the BF muscles displayed one to two bursts of activity during the gait cycle, one in the swing phase and another in the stance phase. This activity pattern aligns with observations in other species for biarticulate muscles such as the BF and semitendinosus muscles in the cat, where the anterior and posterior portions of the muscle may contribute to separate phases of the gait cycle[66], [67].

### Limitations

Minor limitations were identified in this study which did not affect the overall outcomes. First, force plates or pressure mats were not utilized; therefore, the magnitude and patterns of ground reaction forces at different speeds of walking were not quantified. Second, the hoof reflective markers were frequently lost bilaterally during walking, limiting our ability to identify toe-height accurately; nonetheless, the use of the MTP markers provided a very close surrogate. Finally, EMG activity was recorded from only three muscles acting on the hip (abduction and extension) and knee (flexion and extension) in each hindlimb, and no EMG activity was recorded from hip flexors or muscles acting on the ankle. This was because those muscles could not be easily targeted with sEMG. To record activity from the hip flexors or muscles acting on the ankle (e.g., triceps surae and tibialis anterior), percutaneous needle electrodes[68] or implanted electrodes would need to be utilized.

### Conclusions

This study established a baseline of typical quadrupedal locomotion in YMPs. Significant differences in angular kinematics and walking parameters (i.e., stride, swing, and stance duration) were observed as a result of changes in walking speed. Interlimb coordination showed significant changes in both contralateral forelimb-hindlimb and ipsilateral forelimb-hindlimb coordination at increased speed ranges. The muscle activation patterns for the GM, VL, and BF muscles were quantified and only minor differences in onset and offset points were observed with increasing speeds. This study provides normative data against which future studies can detect alterations in the walking metrics of YMPs after SCI and/or neuromodulation in order to assess the effect of interventions on neuroplasticity and functional recovery of locomotion.

## Additional information

### Ethical Statement

All procedures used in this study were approved by the University of Alberta Animal Care and Use Committee.

## References

[1] D. Hirtz, D. J. Thurman, K. Gwinn-Hardy, M. Mohamed, A. R. Chaudhuri, and R. Zalutsky, “How common are the ‘common’ neurologic disorders?,” Neurology, vol. 68, no. 5, pp. 326–337, 2007.

[2] L. D. Hachem, C. S. Ahuja, and M. G. Fehlings, “Assessment and management of acute spinal cord injury: From point of injury to rehabilitation,” J. Spinal Cord Med., vol. 40, no. 6, pp. 665–675, 2017.

[3] K. J. Duberstein et al., “Gait analysis in a pre-and post-ischemic stroke biomedical pig model,” Physiol. Behav., vol. 125, pp. 8–16, 2014.

[4] G. Courtine and M. V. Sofroniew, “Spinal cord repair: advances in biology and technology,” Nat. Med., vol. 25, no. June, 2019, doi: 10.1038/s41591-019-0475-6.

[5] P. J. Reier, M. A. Lane, E. D. Hall, Y. D. Teng, and D. R. Howland, “Translational spinal cord injury research: preclinical guidelines and challenges,” in Handbook of clinical neurology, vol. 109, Elsevier, 2012, pp. 411–433.

[6] J. H. T. Lee et al., “A Novel porcine model of traumatic thoracic spinal cord injury,” J. Neurotrauma, vol. 30, no. 3, pp. 142–159, 2013, doi: 10.1089/neu.2012.2386.

[7] J. T. Hachmann et al., “Large animal model for development of functional Restoration paradigms using epidural and intraspinal stimulation,” PLoS One, vol. 8, no. 12, pp. 1–7, 2013, doi: 10.1371/journal.pone.0081443.

[8] A. N. Dalrymple, D. G. Everaert, D. S. Hu, and V. K. Mushahwar, “A speed-adaptive intraspinal microstimulation controller to restore weight-bearing stepping in a spinal cord hemisection model,” J. Neural Eng., vol. 15, no. 5, 2018, doi: 10.1088/1741-2552/aad872.

[9] J. A. Bamford, R. Marc Lebel, K. Parseyan, and V. K. Mushahwar, “The fabrication, implantation, and stability of Intraspinal Microwire arrays in the spinal cord of cat and rat,” IEEE Trans. Neural Syst. Rehabil. Eng., vol. 25, no. 3, pp. 287–296, 2017, doi: 10.1109/TNSRE.2016.2555959.

[10] R. B. Song, D. M. Basso, R. C. da Costa, L. C. Fisher, X. Mo, and S. A. Moore, “Adaptation of the Basso-Beattie-Bresnahan locomotor rating scale for use in a clinical model of spinal cord injury in dogs,” J. Neurosci. Methods, vol. 268, pp. 117–124, 2016, doi: 10.1016/j.jneumeth.2016.04.023.

[11] S. Wilson et al., “Ovine Hemisection Model of Spinal Cord Injury,” J. Investig. Surg., vol. 0, no. 0, pp. 1–13, 2019, doi: 10.1080/08941939.2019.1639860.

[12] G. Courtine et al., “Kinematic and EMG determinants in quadrupedal locomotion of a non-human primate (Rhesus),” J. Neurophysiol., vol. 93, no. 6, pp. 3127–3145, 2005.

[13] A. Toossi, B. Bergin, M. Marefatallah, B. Parhizi, N. Tyreman, and D. G. Everaert, “Comparative Neuroanatomy of the Lumbosacral Spinal Cord of the Rat , Cat , Pig , Monkey , and Human,” bioRxiv, pp. 1–44, 2020.

[14] K. A. Phillips et al., “Why primate models matter,” Am. J. Primatol., vol. 76, no. 9, pp. 801–827, 2014, doi: 10.1002/ajp.22281.

[15] B. R. Noga, A. J. Santamaria, S. Chang, and D. Francisco, The micropig model of neurosurgery and spinal cord injury in experiments of motor control. Elsevier Inc., 2020.

[16] B. Hoffe and M. R. Holahan, “The Use of Pigs as a Translational Model for Studying Neurodegenerative Diseases,” Front. Physiol., vol. 10, no. July, 2019, doi: 10.3389/fphys.2019.00838.

[17] A. J. Santamaria et al., “Dichotomous Locomotor Recoveries Are Predicted by Acute Changes in Segmental Blood Flow after Thoracic Spinal Contusion Injuries in Pigs,” J. Neurotrauma, vol. 1415, pp. 1399–1415, 2019, doi: 10.1089/neu.2018.6087.

[18] A. Toossi, D. G. Everaert, A. Azar, C. R. Dennison, and V. K. Mushahwar, “Mechanically stable intraspinal microstimulation implants for human translation,” Ann. Biomed. Eng., vol. 45, no. 3, pp. 681–694, 2017.

[19] T. Guiho et al., “Functional Selectivity of Lumbosacral Stimulation : Methodological Approach and Pilot Study to Assess Visceral Function in Pigs,” IEEE Trans. Neural Syst. Rehabil. Eng., vol. 26, no. 11, pp. 2165–2178, 2018, doi: 10.1109/TNSRE.2018.2871763.

[20] Z. Liang et al., “Photobiomodulation by diffusing optical fiber on spinal cord: A feasibility study in piglet model,” J. Biophotonics, vol. 13, no. 4, pp. 1–16, 2020, doi: 10.1002/jbio.201960022.

[21] A. Toossi et al., “Ultrasound-guided spinal stereotactic system for intraspinal implants,” J. Neurosurg. Spine, vol. 29, no. 3, pp. 292–305, 2018.

[22] M. Boakye et al., “Treadmill-Based Gait Kinematics in the Yucatan Mini Pig,” vol. 15, pp. 1–15, 2020, doi: 10.1089/neu.2020.7050.

[23] E. E. Keller et al., “Early sacral neuromodulation ameliorates urinary bladder function and structure in complete spinal cord injury minipigs,” Neurourol. Urodyn., vol. 39, no. 2, pp. 586–593, 2020, doi: 10.1002/nau.24257.

[24] J. Blaszczyk and G. E. Loeb, “Why cats pace on the treadmill,” Physiol. Behav., vol. 53, no. 3, pp. 501–507, 1993, doi: 10.1016/0031-9384(93)90144-5.

[25] G. E. Goslow, R. M. Reinking, and D. G. Stuart, “The cat step cycle: Hind limb joint angles and muscle lengths during unrestrained locomotion,” J. Morphol., vol. 141, no. 1, pp. 1–41, 1973, doi: 10.1002/jmor.1051410102.

[26] M. C. Wetzel, A. E. Atwater, J. V. Wait, and D. G. Stuart, “Neural implications of different profiles between treadmill and overground locomotion timings in cats,” J. Neurophysiol., vol. 38, no. 3, pp. 492–501, 1975, doi: 10.1152/jn.1975.38.3.492.

[27] J. L. Song and J. Hidler, “Biomechanics of overground vs. treadmill walking in healthy individuals,” J. Appl. Physiol., vol. 104, no. 3, pp. 747–755, 2008, doi: 10.1152/japplphysiol.01380.2006.

[28] F. Alton, L. Baldey, S. Caplan, and M. C. Morrissey, “A kinematic comparison of overground and treadmill walking,” Clin. Biomech., vol. 13, no. 6, pp. 434–440, 1998.

[29] D. Tepavac and E. C. Field-Fote, “Vector coding: a technique for quantification of intersegmental coupling in multicyclic behaviors,” J. Appl. Biomech., vol. 17, no. 3, pp. 259–270, 2001.

[30] A. S. P. Varejao and V. M. Filipe, “Contribution of cutaneous inputs from the hindpaw to the control of locomotion in rats,” Behav. Brain Res., vol. 176, no. 2, pp. 193–201, 2007.

[31] G. C. Koopmans et al., “Strain and locomotor speed affect over-ground locomotion in intact rats,” Physiol. Behav., vol. 92, no. 5, pp. 993–1001, 2007, doi: 10.1016/j.physbeh.2007.07.018.

[32] A. R. Wu, C. S. Simpson, E. H. F. van Asseldonk, H. van der Kooij, and A. J. Ijspeert, “Mechanics of very slow human walking,” Sci. Rep., vol. 9, no. 1, pp. 1–10, 2019.

[33] G. E. Goslow Jr, R. M. Reinking, and D. G. Stuart, “The cat step cycle: hind limb joint angles and muscle lengths during unrestrained locomotion,” J. Morphol., vol. 141, no. 1, pp. 1–41, 1973.

[34] J.-P. Gossard et al., “The spinal generation of phases and cycle duration,” Prog. Brain Res., vol. 188, pp. 15–29, 2011.

[35] A. Frigon, “Central pattern generators of the mammalian spinal cord,” Neuroscientist, vol. 18, no. 1. pp. 56–69, 2012, doi: 10.1177/1073858410396101.

[36] M. C. Wetzel, A. E. Atwater, J. V Wait, and D. C. Stuart, “Neural implications of different profiles between treadmill and overground locomotion timings in cats,” J. Neurophysiol., vol. 38, no. 3, pp. 492–501, 1975.

[37] D. Wisleder, R. F. Zernicke, and J. L. Smith, “During the Swing Phase of Cat Locomotion,” Exp. Brain Res., pp. 651–660, 1990.

[38] A. W. English, “Interlimb coordination during stepping in the cat: The role of the dorsal spinocerebellar tract,” Exp. Neurol., vol. 87, no. 1, pp. 96–108, 1985, doi: 10.1016/0014-4886(85)90136-0.

[39] S. K. Patrick, J. A. Noah, and J. F. Yang, “Interlimb coordination in human crawling reveals similarities in development and neural control with quadrupeds,” J. Neurophysiol., vol. 101, no. 2, pp. 603–613, 2009, doi: 10.1152/jn.91125.2008.

[40] L. Righetti, A. Nylén, K. Rosander, and A. J. Ijspeert, “Kinematic and gait similarities between crawling human infants and other quadruped mammals,” Front. Neurol., vol. 6, no. FEB, pp. 1–11, 2015, doi: 10.3389/fneur.2015.00017.

[41] Hildebra.M, “Symmetrical Gaits of Primates,” Am. J. Phys. Anthropol., vol. 26, no. 2, pp. 119-, 1967.

[42] T. Warabi, M. Kato, K. Kiriyama, T. Yoshida, and N. Kobayashi, “Treadmill walking and overground walking of human subjects compared by recording sole-floor reaction force,” Neurosci. Res., vol. 53, no. 3, pp. 343–348, 2005, doi: 10.1016/j.neures.2005.08.005.

[43] T. Jung, Y. Kim, L. E. Kelly, M. Wagatsuma, Y. Jung, and M. F. Abel, “Comparison of Treadmill and Overground Walking in Children and Adolescents,” Percept. Mot. Skills, vol. 128, no. 3, pp. 988–1001, 2021.

[44] A. Frigon, “The neural control of interlimb coordination during mammalian locomotion,” J. Neurophysiol., vol. 117, no. 6, pp. 2224–2241, 2017, doi: 10.1152/jn.00978.2016.

[45] S. Grillner, “Control of Locomotion in Bipeds, Tetrapods, and Fish,” Compr. Physiol., 2011, doi: 10.1002/cphy.cp010226.

[46] A. Frigon, M.-F. Hurteau, Y. Thibaudier, H. Leblond, A. Telonio, and G. D’Angelo, “Split-belt walking alters the relationship between locomotor phases and cycle duration across speeds in intact and chronic spinalized adult cats,” J. Neurosci., vol. 33, no. 19, pp. 8559–8566, 2013.

[47] C. Langlet, H. Leblond, and S. Rossignol, “Mid-lumbar segments are needed for the expression of locomotion in chronic spinal cats,” J. Neurophysiol., vol. 93, no. 5, pp. 2474–2488, 2005, doi: 10.1152/jn.00909.2004.

[48] K. E. Miller, V. D. Douglas, A. B. Richards, M. J. Chandler, and R. D. Foreman, “Propriospinal neurons in the C1-C2 spinal segments project to the L5-S1 segments of the rat spinal cord,” Brain Res. Bull., vol. 47, no. 1, pp. 43–47, 1998.

[49] S. Miller, D. J. Reitsma, and F. G. A. Van Der Meche, “Functional organization of long ascending propriospinal pathways linking lumbo-sacral and cervical segments in the cat,” Brain Res., vol. 62, no. 1, pp. 169–188, 1973.

[50] K. G. Pearson, “Generating the walking gait: role of sensory feedback,” Prog. Brain Res., vol. 143, pp. 123–129, 2004.

[51] A. Frigon and S. Rossignol, “Experiments and models of sensorimotor interactions during locomotion,” Biol. Cybern., vol. 95, no. 6, pp. 607–627, 2006.

[52] A. W. English, “Interlimb coordination during stepping in the cat: An electromyographic analysis,” J. Neurophysiol., vol. 42, no. 1, pp. 229–243, 1979, doi: 10.1152/jn.1979.42.1.229.

[53] A. Goswami, “A new gait parameterization technique by means of cyclogram moments: Application to human slope walking,” Gait Posture, vol. 8, no. 1, pp. 15–36, 1998.

[54] J. Charteris, D. Leach, and C. Taves, “Comparative kinematic analysis of bipedal and quadrupedal locomotion: a cyclographic technique.,” J. Anat., vol. 128, no. Pt 4, p. 803, 1979.

[55] Y. Higurashi et al., “Locomotor kinematics and EMG activity during quadrupedal versus bipedal gait in the Japanese macaque,” J. Neurophysiol., vol. 122, no. 1, pp. 398–412, 2019.

[56] C. Hershler and M. Milner, “Angle--angle diagrams in the assessment of locomotion.,” Am. J. Phys. Med., vol. 59, no. 3, pp. 109–125, 1980.

[57] E. C. Field-Fote and D. Tepavac, “Improved intralimb coordination in people with incomplete spinal cord injury following training with body weight support and electrical stimulation,” Phys. Ther., vol. 82, no. 7, pp. 707–715, 2002.

[58] M. Iosa, L. Gizzi, F. Tamburella, and N. Dominici, “Neuro-motor control and feed-forward models of locomotion in humans,” Front. Hum. Neurosci., vol. 9, p. 306, 2015.

[59] M. Philippson, L’autonomie et la centralisation dans le système nerveux des animaux. Falk, 1905.

[60] Y. P. Ivanenko, R. E. Poppele, and F. Lacquaniti, “Five basic muscle activation patterns account for muscle activity during human locomotion,” J. Physiol., vol. 556, no. 1, pp. 267–282, 2004.

[61] M. Belanger, T. Drew, J. Provencher, and S. Rossignol, “A comparison of treadmill locomotion in adult cats before and after spinal transection,” J. Neurophysiol., vol. 76, no. 1, pp. 471–491, 1996.

[62] F. Lacquaniti, Y. P. Ivanenko, A. d’Avella, K. Zelik, and M. Zago, “Evolutionary and developmental modules,” Front. Comput. Neurosci., vol. 7, p. 61, 2013.

[63] G. Catavitello, Y. Ivanenko, and F. Lacquaniti, “A kinematic synergy for terrestrial locomotion shared by mammals and birds,” Elife, vol. 7, p. e38190, 2018.

[64] S. Grillner and A. El Manira, “Current principles of motor control, with special reference to vertebrate locomotion,” Physiol. Rev., vol. 100, no. 1, pp. 271–320, 2020.

[65] N. Dominici et al., “Locomotor Primitives in Newborn Babies and Their Development,” vol. 334, no. November, pp. 997–1000, 2011.

[66] A. W. M. English and O. I. Weeks, “An anatomical and functional analysis of cat biceps femoris and semitendinosus muscles,” J. Morphol., vol. 191, no. 2, pp. 161–175, 1987.

[67] J. L. Smith, S. H. Chung, and R. F. Zernicke, “Gait-related motor patterns and hindlimb kinetics for the cat trot and gallop,” Exp. brain Res., vol. 94, no. 2, pp. 308–322, 1993.

[68] A. Toossi et al., “Effect of anesthesia on motor responses evoked by spinal neural prostheses during intraoperative procedures,” J. Neural Eng., vol. 16, no. 3, p. 36003, 2019.

